# A window of cell cycle plasticity enables imperfect regeneration of an adult postmitotic organ in *Drosophila*

**DOI:** 10.1101/2025.06.05.658044

**Authors:** Navyashree A. Ramesh, Arpitha S. Banakar, Isabella Karanja, Robin E. Harris, Laura A. Buttitta

## Abstract

The *Drosophila* ejaculatory duct (ED) is a secretory tissue of the male somatic reproductive system responsible for producing components of the seminal fluid which support fertility, serve antimicrobial functions and influence the physiological changes in the female after mating. The ED is a simple organ made up of secretory epithelial cells that are encased by extracellular matrix and a layer of innervated contractile muscle. The ED secretory epithelial cells are post-mitotic and lack known stem cells or progenitors in the adult, but they are not fully quiescent. They undergo a variant cell cycle called endoreplication immediately post-eclosion to increase organ size and protein synthesis capacity. Polyploid and post-mitotic tissues often face unique challenges in response to cell loss due to their inability to proliferate. Here, we show that the adult ED is capable of significant recovery after cell loss due to a combination of increased nuclear and cellular hypertrophy that partially restores tissue mass and organ function. The early cell cycle plasticity of this adult tissue is critical for this recovery, as older tissues that have few or no endocycles exhibit reduced capacity for recovery after cell loss. Together, our findings establish the *Drosophila* ED as a model to study post-mitotic polyploid tissue repair and highlight a combination of endocycles and hypertrophy as a key mechanism for functional regeneration in the absence of mitosis.

## Introduction

The *Drosophila* male ejaculatory duct (ED) serves as conduit for sperm and seminal fluid transfer to the female via the ejaculatory bulb during mating. In addition to its transport function, the ED itself is a secretory organ, producing a substantial number of seminal fluid proteins (SFPs) including antimicrobial peptides and peptides involved in a range of reproductive functions (Date-Ito et al., 2002; Imamura et al., 1998; Lung et al., 2001; Richmond et al., 1980; Sturm et al., 2021; Takemori and Yamamoto, 2009). Anatomically, the ED is composed of a basal, innervated muscular layer and an apical layer of polyploid secretory cells (Ramesh et al., 2025; Susic-Jung et al., 2012). These secretory cells are the primary source of ED-derived fertility factors.

ED secretory cells of newly eclosed adult are post-mitotic, but not fully quiescent (Ramesh et al., 2025). Hours after eclosion, secretory cells of the ED undergo a variant cell cycle known as endocycling (Edgar et al., 2014). Endocycling cells alternate between G1 and S phases without undergoing mitosis, resulting in polyploidy (Ramesh et al., 2025). The post eclosion endocycles are essential in the secretory cells for post-eclosion growth of the ED and full male fertility (Ramesh et al., 2025). Endocycles in the ED are rare after 48h and cease by 72h post-eclosion, after which the cells remain fully quiescent. This tissue contains no known stem or progenitor cell population and thus is thought to have very limited to no regenerative capability.

Tissues respond to cell loss by initiating regenerative programs to restore lost mass and function. In mitotically active tissues, this typically involves compensatory cellular proliferation (CCP), where either resident stem cells or surrounding progenitor cells divide to replace the lost cells (de Morree and Rando, 2023; Xia et al., 2018). However, largely post-mitotic tissues such as the heart, kidneys, cornea, bladder, and brain possess relatively few or no regenerative stem cell populations (Lavecchia et al., 2022; Losick, 2016; Shu et al., 2018). Understanding how postmitotic tissues respond to injury, to maintain structural integrity and partially restore function remains a major challenge in regenerative biology.

Here, we show that the secretory cells of the young adult male ED retain a surprising capacity to restore tissue morphology and recover fertility function after genetically-driven partial organ ablation. In response to cell loss, the window for endocycling in the young adult ED is extended temporarily, providing an opportunity for postmitotic polyploid cells to restore tissue mass in part through compensatory endocycles. As the tissue matures and cell cycle plasticity is lost, the regenerative capacity of this tissues declines, suggesting that the regulators of endocycle exit and entry into quiescence limit this organ’s recovery after damage. We propose the ED can be a genetically tractable model to study postmitotic regeneration and cell cycle plasticity in a terminally differentiated adult tissue.

## Results & Discussion

### Apoptotic cell loss in the ED occurs through apical extrusion and leads to irreversible cell loss

Secretory cells of the ED in newly eclosed males undergo a variant cell cycle called endoreplication, in which they replicate their DNA without entering mitosis, thereby increasing cellular ploidy. We previously showed that post-eclosion endocycles in this tissue are essential for full male fertility (Ramesh et al., 2025). However direct studies of secretory cell loss in the ED have not been performed. We induced apoptosis in the secretory cells of the adult ED by expressing the IAP antagonist *Reaper* (Ryoo et al., 2002; Sandu et al., 2010) using an ED-secretory cell specific *Luna-Gal4* (Ramesh et al., 2025) (Figure 1A). Inducing *Reaper* in the ED led to cells with pyknotic nuclei, indicating that the cells were undergoing apoptosis (Figure 1B). By co-expressing RFP tagged with a nuclear localization signal (*UAS-nls-RFP*) we could observe cell loss (Suppl Fig 1A), and disruption of epithelial cell-cell contacts, such as septate junctions labeled with Fasciclin III (Fas III). Following nearly complete ablation of secretory cells, only muscle/neuronal nuclei, which were Elav-positive, remained in the ED (Suppl Fig 1A).

**Figure 1.**
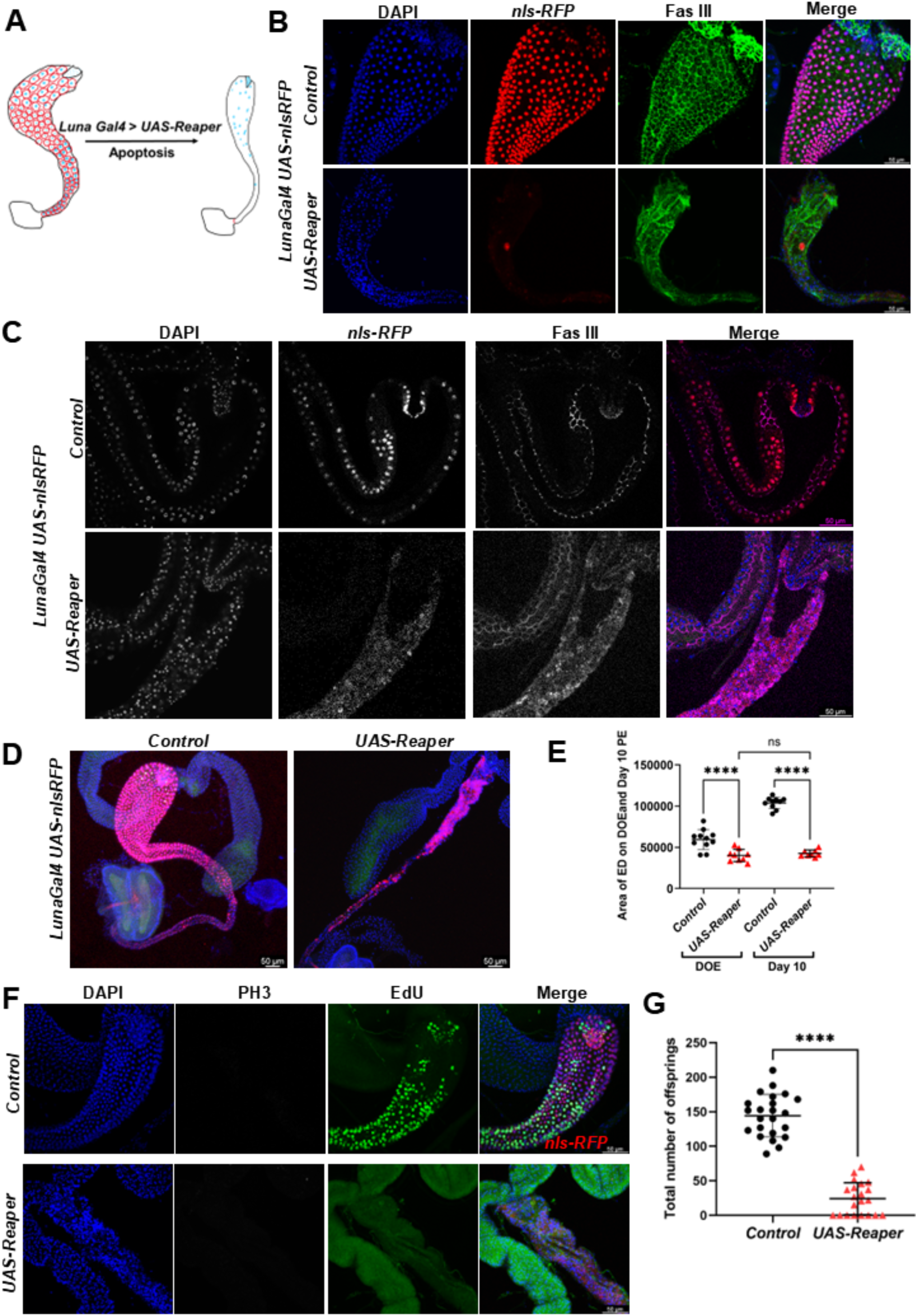
Reaper expression induces apoptosis in *Drosophila* ED secretory cells. (A) Schematic showing targeted expression of Reaper in ED. (B) Confocal images of control (Luna Gal4, UAS-RFP without UAS-Reaper), and constitutive Reaper (Luna Gal4, UAS-RFP + UAS-Reaper) at Day 10 post-eclosion (PE), immunolabeled with FasIII, DAPI. (C) Single optical sections of EDs through the lumen on the day of eclosion (DOE) stained with FasIII. (D) Confocal image of the male reproductive tract, highlighting the reduced size of ED upon persistent Reaper expression. (E) Quantification of control ED vs Reaper-expressing ED area on Day 0 and Day 10 PE. One-way ANOVA with multiple comparisons. P<0.0001****. (F) Confocal images of EDs stained with EdU, PH3 and DAPI. Unpaired two-tailed t-test. P<0.0001****. (G) Quantification of male fertility in control and Reaper-expressing animals. Scale bar:50μm.

Upon cell death, the ED secretory cells underwent apical extrusion into the ED lumen (Figure 1C). These apoptotic cells were further expelled through the ejaculatory bulb (Figure 1B). Despite the loss of most secretory cells, the external basal muscle layer remained largely unaffected (Supplementary Figure 1C), while organ size was significantly reduced (Figure 1D,E). To determine whether epithelial cell loss triggered proliferation or cell cycles within the diploid cells of the muscle layer or adjacent non-Luna Gal4 expressing tissues, such as the diploid anterior ED papillae, we fed the animals 5-Ethynyl-2′-deoxyuridine (EdU) to label S-phases and stained for phospho-Histone H3 (PH3 Ser10) to visualize mitotic activity. We observed no evidence of cell cycles in the muscle layer or nearby ED papillae in response to the loss of the secretory cells (Figure 1F). We also did not observe any evidence of multinucleated cells that would result from cell fusion.

Our previous study revealed that blocking post-eclosion endoreplication in ED secretory cells led to reduced fertility and fewer offspring (Ramesh et al., 2025). In line with this, loss of ED secretory cells severely affected fertility (Figure 1G). We used quantitative video assays to ensure that ablation of the ED secretory cells did not affect male mating behaviors, and the mating attempts by males with ED damage were performed at the same frequency and rate as control males without tissue damage. Post fertility assay, we dissected the male EDs and observed that sperm becomes entrapped within the ED lumen when the ED secretory layer is severely disrupted. By contrast, undamaged mated males very rarely retain significant amounts of sperm in the duct (Supplementary Figure 1D). It is likely that the ED cells secrete factors required for the effective transfer of sperm, seminal fluid proteins, and other peptides to the female reproductive tract.

### Recovery from transient damage in the anterior region of the Ejaculatory Duct

To investigate whether the secretory epithelium can recover following transient cell loss, we employed a conditional approach using the *Gal4/UAS-Reaper* system in combination with a temperature-sensitive *Gal80* repressor (*Gal80^ts^*). At a lower temperature (18 °C), Gal80 is active and represses Gal4-mediated transcription (Figure 2A). However, at higher temperatures (25–29°C), Gal80^ts^ becomes inactive, allowing Gal4 to drive transgene expression (McGuire et al., 2004). To validate the *Gal80^ts^* system with the ED driver, we raised flies carrying *Luna-Gal4, UAS-nls-RFP*, and *Gal80^ts^* at 18 °C. At this temperature, *Gal80^ts^* repressed Gal4, and no *nls-RFP* expression was detected. Upon shifting the flies to 29 °C, we observed robust *nls-RFP* expression in the ED (Suppl Fig 2A), indicating activation of *Gal4* due to *Gal80^ts^* inactivation. To determine how quickly Gal4 activity is repressed after returning to 18 °C, we used a *destabilized GFP (dsGFP)* reporter, which is rapidly degraded in the absence of *Gal4* activity, providing a readout of transient gene expression. Flies expressing *Luna-Gal4* and *UAS-dsGFP*, were maintained at 18 °C and showed no GFP signal. After shifting to 29 °C, GFP expression appeared in the anterior region of the ED after 32 hours, with no expression in the posterior ED. We then shifted the animals back to 18 °C to assess the time required for Gal4 repression following *Gal80^ts^* reactivation. We observed that within 24 hours, GFP fluorescence was completely lost, indicating detectable repression of Gal4 and degradation of *dsGFP* (Suppl Fig 2B).

**Figure 2.**
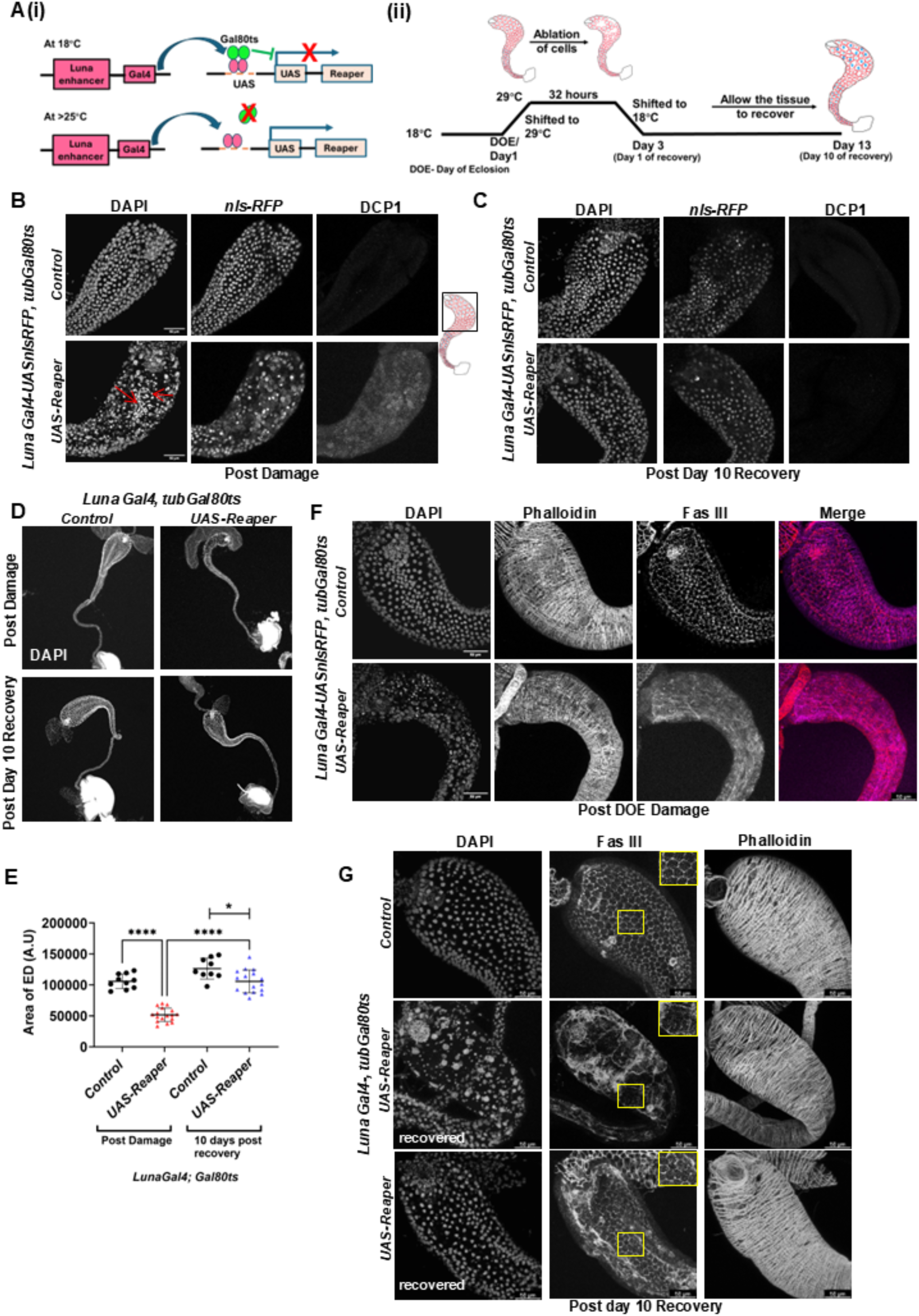
Recovery from transient damage in the anterior region of the Ejaculatory Duct. (A)(i) Schematic of the temperature-sensitive Gal80^ts^ system used to transiently induce Luna-Gal4-mediated expression of Reaper in ED cells. (ii) Illustration of ED damage and recovery paradigm: Flies are housed at 18°C to suppress Gal4 via Gal80^ts^. On the day of eclosion, flies are shifted to 29°C for 32 hours to induce Reaper, leading to apoptosis of secretory cells in the anterior ED. Flies are then returned to 18°C for 10 days to allow for tissue recovery. (B) Confocal images of the anterior region of ED immediately after 32 hours of damage induction at 29°C. Tissues are stained with DCP-1 and DAPI. (C) Confocal images of anterior ED post day10 recovery, stained with DCP-1, DAPI. (D) Confocal images of whole EDs post-damage and post day10 recovery (10×magnification). (E) Quantification of ED area. One-way ANOVA with multiple comparisons. P<0.0001****. (F) Confocal images of anterior ED region post 32 hours damage stained with Phalloidin, FasIII and DAPI. (G) Confocal images of anterior ED after 10-days of recovery stained with Phalloidin, FasIII, and DAPI. The middle row represents partial recovery, with cellular hypertrophy. The bottom panel shows and example of full recovery of ED morphology. Scale bar:50μm.

Based on these observations, we shifted *Luna-Gal4/UAS-ReaperGal80^ts^* animals from 18 °C to 29 °C on the day of eclosion (DOE) for 32 hours to induce transient damage (Figure 2A). As the ED is a post-mitotic tissue with secretory cells that undergo endoreplication for the first 2 days post-eclosion, we chose to induce damage on the day of eclosion, during the window of endogenous endoreplication. Upon activation of *Reaper*, we observed caspase-induced apoptosis in cells located in the anterior ED, marked by the presence of multiple pyknotic nuclei (Figure 2B). In contrast, no pyknotic nuclei were observed in the posterior region within 32h of *Reaper* induction (Suppl Figure 2D), indicating region-specific cell ablation. We observed a similar pattern with an antibody against cleaved Death caspase-1 (DCP-1, Suppl Figure 2D) as well as a *Drice*-Based Sensor GFP (*DBS-GFP*) which indicate active caspase-mediated cleavage of substrates (Baena-Lopez et al., 2018) (Suppl Figure 2C, 2E). The anterior ED contains larger secretory cells with higher ploidy levels of 16-32C, while the middle and posterior ED contains smaller secretory cells with lower ploidies of 8-4C. We speculate that the increased Gal4 copy number and expression level in the anterior ED preferentially induces apoptosis with this limited activation window.

With transient *UAS-Reaper* expression, 100% of animals exhibited loss of secretory cells in the anterior region of the ejaculatory duct (Supplementary Figure 2F). Cell loss could be completely suppressed by co-expression of the baculoviral apoptosis inhibitor *UAS-P35* with *UAS-Reaper* (Hay et al., 1994). Interestingly, organs co-expressing *UAS-P35* with *UAS-Reaper* appeared indistinguishable from control animals without Reaper, suggesting that apoptosis-induced proliferation does not take place in this tissue (Suppl Figure 3) (Perez-Garijo et al., 2004; Perez-Garijo et al., 2009). Thus, our system efficiently induces partial ablation in the anterior ED through caspase-induced apoptosis.

We shifted the animals back to 18 °C to repress *Reaper* expression via *Gal80^ts^* and maintained them at this temperature for 10 days to observe tissue response to damage (Figure 2A). Remarkably, we observed substantial recovery of the ED 10 days post-damage. Cells were no longer DCP1-positive, indicating the resolution of apoptosis (Figure 2C and Suppl Figure 2E). In the animals after a 10-day recovery without Reaper expression, we observed near-normal tissue size (Figure 2D). As our damage paradigm induces cell loss in 100% of animals, we interpret this to be evidence of significant restoration of both organ mass and epithelial integrity in this postmitotic and terminally differentiated tissue (Figure 2E).

### The ED undergoes compensatory cellular hypertrophy (CCH) in response to cell loss

The loss of anterior secretory cells led to organ shrinkage, however we observed no disruption of the muscle following secretory cell loss. By contrast, epithelial septate junctions were severely disrupted immediately after damage (Figure 2F), but after 10 days of recovery at 18 °C septate junctions were restored and appeared normal (Figure 2G). We observe cell loss in 100% of animals immediately after damage, but the degree of cell loss is variable. In cases where cell loss appears mild, we observe enlarged epithelial cells with normal or near normal nuclear sizes, suggestive of cellular growth or stretching without increased ploidy (Figure 2G, bottom row). However in cases where many secretory cells are lost, the remaining cells undergo compensatory cellular hypertrophy (CCH) with visibly enlarged nuclei, indicative of an increase in cellular ploidy (Tamori and Deng, 2013, 2014) (Figure 2G, middle row). Cells are significantly larger after damage followed by recovery compared to undamaged controls (Figure 3A) with enlarged nuclei (Figure 3B). Thus, the secretory cells of the ED are capable of growth after mild cell loss and compensatory cellular hypertrophy (CCH) in response to more severe cell loss to restore lost tissue mass.

**Figure 3.**
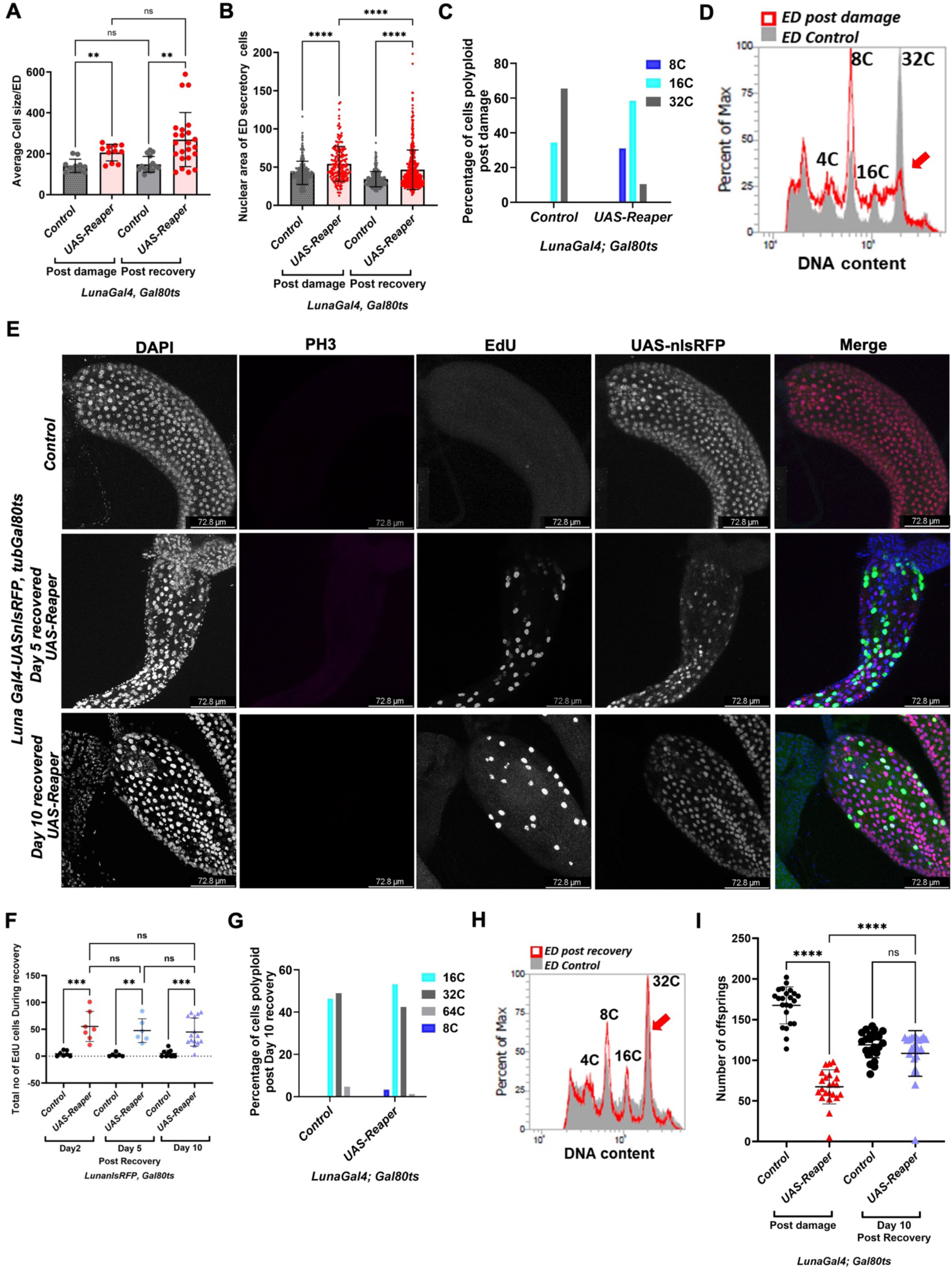
Endoreplication in the ED after cell loss restores ploidy, tissue mass and supports functional regeneration. (A) Quantification of average secretory cell size per ED. Each dot represents the mean cell size per animal. (B) Quantification of nuclear area of ED secretory cells in anterior region. Each dot represents the area of an individual nucleus. (C) Quantification of ploidy in secretory cells of the anterior ED, measured by DAPI intensity, after 32 hours damage (D) Flow cytometric analysis of nuclear ploidy in control and Reaper-expressing EDs post 32 hours damage. Grey peak-control ED nuclei, red peak nuclei from Reaper EDs. Red arrow indicates the drop in 32C nuclei after Reaper induced damage. (E) Confocal micrographs of the anterior ED Control, Day5 recovered, and Day10 recovered stained for EdU, PH3. (F) Quantification of EdU-positive cells in control and Reaper EDs from images of (E). (G) Quantification of ploidy in the anterior secretory cells of control and Reaper EDs post day10 recovery. (H) Flow cytometric analysis of nuclear ploidy in control and Reaper-expressing EDs after Day10 of recovery. Red arrow indicates the restoration of the 32C population after 10 days of recovery. (I) Quantification of male fertility in control and Reaper-expressing animals. Damaged ED: Mating performed post 32hours damage at 29°. Recovered ED: Mating performed day10 post-recovery at 18°C. Scale bar:50μm. Statistical analysis: One-way ANOVA with multiple comparisons. P<0.0001 ****.

To quantify the ploidy of the cells in the anterior region, we measured the integrated DAPI intensity from nuclei, normalized to diploid cells of the innervated muscle within the same organ. The ploidy of the ED immediately post-damage was significantly reduced compared to controls (Figure 3C), consistent with loss of the anterior-most, highest ploidy ED cells. This increased the number of 8C and 16C cells in the anterior region immediately post-damage (compared to 32C in controls). By contrast, after 10 days of recovery at 18°C, we found that cellular ploidies in the anterior region were nearly fully restored (Figure 3G), with the reappearance of many 32C cells in the anterior ED. We confirmed these results using flow cytometry, showing loss of the highest ploidy cells immediately after damage (Figure 3D, red arrow) with substantial restoration of 32C cells after recovery (Figure 3H, red arrow). Notably the effect on ploidy in the flow cytometry assay is muted compared to the imaging quantifications because the flow assay uses nuclei isolated from the entire ED, while our imaging ploidy measurements reveal differences that are specific to the anterior region of the ED, where the most severe cell loss occurs. As a whole, these findings support that an increase in cellular ploidy accompanies the observed CCH following damage and recovery.

To visualize DNA synthesis during CCH, we fed the animals EdU immediately after the 32-hour damage period and tissues were dissected after 2, 5 and 10 days of recovery in the presence of EdU. We co-stained for phospho-Histone H3 (PH3 Ser10) to detect any mitotic activity. At Day 5, we observed many EdU-positive cells (Figure 3E, middle panel), however, no PH3-positive cells were detected throughout the secretory and muscle layers of the ED. Control undamaged EDs (Figure 3E, top panel) showed no EdU-positive cells, consistent with our prior observations that the tissue completes endocycling by Day 3 post-eclosion normally (Ramesh et al., 2025). We included *Drosophila* larval imaginal discs as a positive control for PH3 staining, which undergo robust mitotic proliferation (Figure 3E, Suppl Figure 4B). By Day 5, despite significant EdU labeling, the anterior ED remained smaller than controls, but by Day 10, the ED tissue mass was fully restored, retained significant EdU labeling, and exhibited no evidence of mitoses. This time course indicates that most endoreplication likely occurs between Days 2 - 4 of recovery (Figure 3F). This is equivalent to ∼72-96 hours post-eclosion without damage, a time when the ED has normally exited the endocycle and is fully quiescent (Ramesh et al., 2025). This suggests that following secretory cell ablation during the post-eclosion endocycle, the remaining ED cells extend their endocycling window to complete additional damage-induced endocycles as a compensatory mechanism. Following the compensatory endoreplication, the tissue eventually regains mass through increased cell growth. We observed similar effects using transient induction of the apoptotic pathway component Hid, confirming these effects on the endocycle in surviving ED cells are likely in response to apoptotic cell loss (Suppl. Fig. 4).

To test whether secretory cell restoration in the ED could originate from other cell types in or near the ED, we performed G-TRACE labeling after induced damage using the *Luna-Gal*4 driver. G-TRACE (Gal4 Technique for Real-time and Clonal Expression) enables visualization of both real-time (RFP) and permanent Gal4-lineage driven expression (GFP) (Evans et al., 2009). The *Luna-Gal4* driver used here is specific to the secretory cells of the ED in the male reproductive system and is active in the ED secretory cells from 1 day prior to eclosion throughout adulthood (Ramesh et al., 2025). We found that all secretory cells in the ED co-express both GFP and RFP, immediately after damage, as well as 10 days post-damage (Suppl Figure 5E). These findings suggest that tissue mass recovery for the ED occurs from within the preexisting Luna-Gal4-expressing lineage of differentiated secretory cells, without the involvement of stem or progenitor cells.

Perfect regeneration restores tissue mass, patterning and function after cell loss. Tissues with few or no progenitors or stem cells may undergo “imperfect” regenerative mechanisms which may fail to restore cell numbers and prior tissue pattern, but can aid in restoring tissue or organ function, as in the mammalian liver (Cordero-Espinoza and Huch, 2018; Poss and Tanaka, 2024). To test whether the CCH we observe in the ED restores male fertility, we performed fertility assays on males during damage as well as after recovery. In brief, we crossed males expressing *UAS-Reaper* to Canton-S virgin females at a male-to-female ratio of 1:8, either at 29 °C to sustain Reaper-induced damage during mating or after 10 days of recovery at 18 °C. Males with damaged EDs produced significantly fewer offspring (Figure 3I), and exhibited sperm trapped within the ED (Suppl Figure 5C). By contrast, males that recovered after transient damage for 10 days exhibited significantly improved fertility (Figure 4D) and lacked sperm entrapped within the ED lumen, indicating that the tissue had recovered function. Thus, the post-mitotic ED is capable of partial functional regeneration following transient loss of anterior secretory cells.

**Figure 4.**
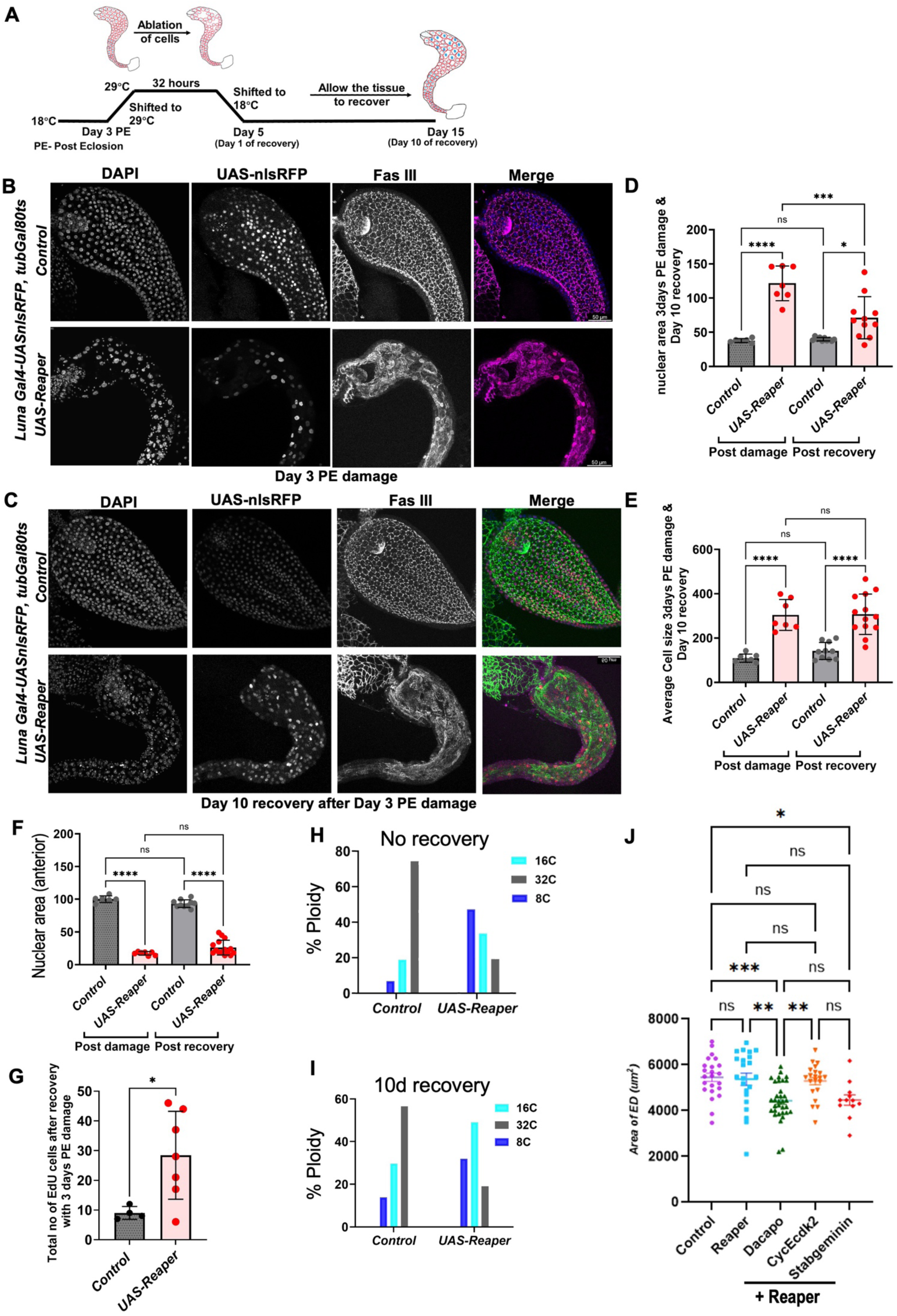
Cell cycle plasticity is required for ED regeneration. (A) Schematic of damage and recovery paradigm for Day3 post-eclosion damage induction. Flies housed at 18°C suppress Gal4 activity via Gal80^ts^. On Day3 post-eclosion, flies were shifted to 29°C for 32 hours to induce Reaper expression. Flies were returned to 18°C for 10 days to allow recovery. (B) Confocal images of anterior ED region immediately after damage on Day3 stained with FasIII, DAPI. (C) Confocal images of anterior ED region after 10 days of recovery at 18°C stained with FasIII, DAPI. (D) Quantification of nuclear area of secretory cells after damage on Day3 and following 10 days of recovery. Each dot represents average nuclear area per ED. One-way ANOVA with multiple comparisons. P<0.0001****. (E) Quantification of average cell size of secretory cells after damage on Day3 and following 10 days of recovery. (F) Quantification of total nuclei per unit area in anterior ED post Day3 damage and recovery. (G) Quantification of EdU-positive secretory cells in undamaged controls and after damage on Day3 following 10 days of recovery in the presence of EdU. Unpaired two-tailed t-test. P<0.0001****. (H) Quantification of ploidy in anterior secretory cells after damage on Day3. (I) Quantification of ploidy in anterior secretory cells after damage on Day3 followed by 10 days of recovery. (J) Quantification of ED area for the indicated genotypes after 10 days of recovery. One-way ANOVA with multiple comparisons. P<0.0001****. Scale bar:50μm.

### Cell cycle plasticity is required for ED regeneration

Thus far, our damage induction paradigm begins on the day of eclosion, when ED secretory cells are still actively undergoing endocycling. However, this window of cell cycle plasticity closes by 48-72h post eclosion, when nearly all secretory ED cells exit the endocycle and become quiescent. To test whether active endocycling is required for ED recovery after damage, we induced ED cell death on Day 3 post-eclosion by shifting the males to 29 °C for 32 hours to activate Reaper expression (Figure 4A). We observed ablation of secretory cells from the anterior ED, with noticeable shrinkage of the anterior region compared to controls (Figure 4B). This loss of cells also disrupted epithelial cell-cell junctions, as indicated by Fas III staining. We then shifted the damaged males back to 18 °C for 10 days to allow for recovery. However, the tissue was unable to exhibit the level of recovery we observed when damage was induced on the day of eclosion. The anterior ED remained smaller, and although some compensatory cellular hypertrophy (CCH) was observed, cell junctions were not fully restored (Figure 4C). Even after 10 days, the tissue remained small, with reduced tissue mass. We saw no significant difference in cell size or recovery of cell density after 10 days of recovery (Figure 4E, 4F).

We also examined EdU labeling in males damaged at either day 3 or day 5 followed by EdU feeding during 10 days of recovery. We observed increased EdU labeling in animals damaged at day 3 compared to undamaged controls (Figure 4G). However, the number of EdU-positive cells in response to damage induced on day 3 was substantially lower than that observed after damage on the day of eclosion, and was further reduced when damage was induced on day 5. This suggests that the window of endocycling can be extended to day 3, but by day 5 ED cells exhibit very low capacity to re-enter the endocycle thereby limiting tissue recovery. Consistent with this, we observed a loss of higher-ploidy (32C) cells immediately after damage (Figure 4H), and these high-ploidy cells did not recover, even after 10 days (Figure 4I).

To further confirm the role of the endocycle in promoting post damage organ recovery, we induced damage using Reaper immediately upon eclosion for 32h in the presence of positive and negative cell cycle regulators to enhance or inhibit the capacity for the ED secretory cells to endocycle and shifted back to 18°C for 10d of recovery. We observed that transient induction of the G1-S Cyclin/Cdk CyclinE during damage had little effect on organ size after recovery, while transient induction of the G1-S Cyclin/Cdk inhibitor Dacapo and a stabilized form of the DNA replication licensing inhibitor Geminin significantly reduced organ size recovery (Fig 4J). This demonstrates that reducing the capacity to enter S-phase after damage limits organ recovery. Together, these results suggest that the early cell cycle plasticity allowed by an endoreplication-competent window during organ maturation is likely to be essential for the regenerative capacity of post-mitotic ED secretory cells following transient damage.

### Necrotic cell loss induces compensatory cellular hypertrophy (CCH) and endocycling in the ED

We next examined whether the ED could recover from another form of transient cell loss, induced necrosis. We induced necrosis in the secretory epithelium using inducible expression of *UAS*-GluR1, a murine glutamate receptor, which we and others previously showed causes cellular Ca^2+^ influx leading to osmotic swelling and necrosis (Klemm et al., 2021; Yang et al., 2013). Persistent expression of GluR1 for 10 days caused extensive cell loss in the ED and led to induction of the apoptotic marker, cleaved Dcp1 (Fig. 5A-C). This demonstrates that a phenomenon we previously described in proliferating wing discs, necrosis-induced caspase positivity (Klemm et al., 2021; Klemm et al., 2025), also can occur in postmitotic tissues experiencing prolonged necrosis.

**Figure 5.**
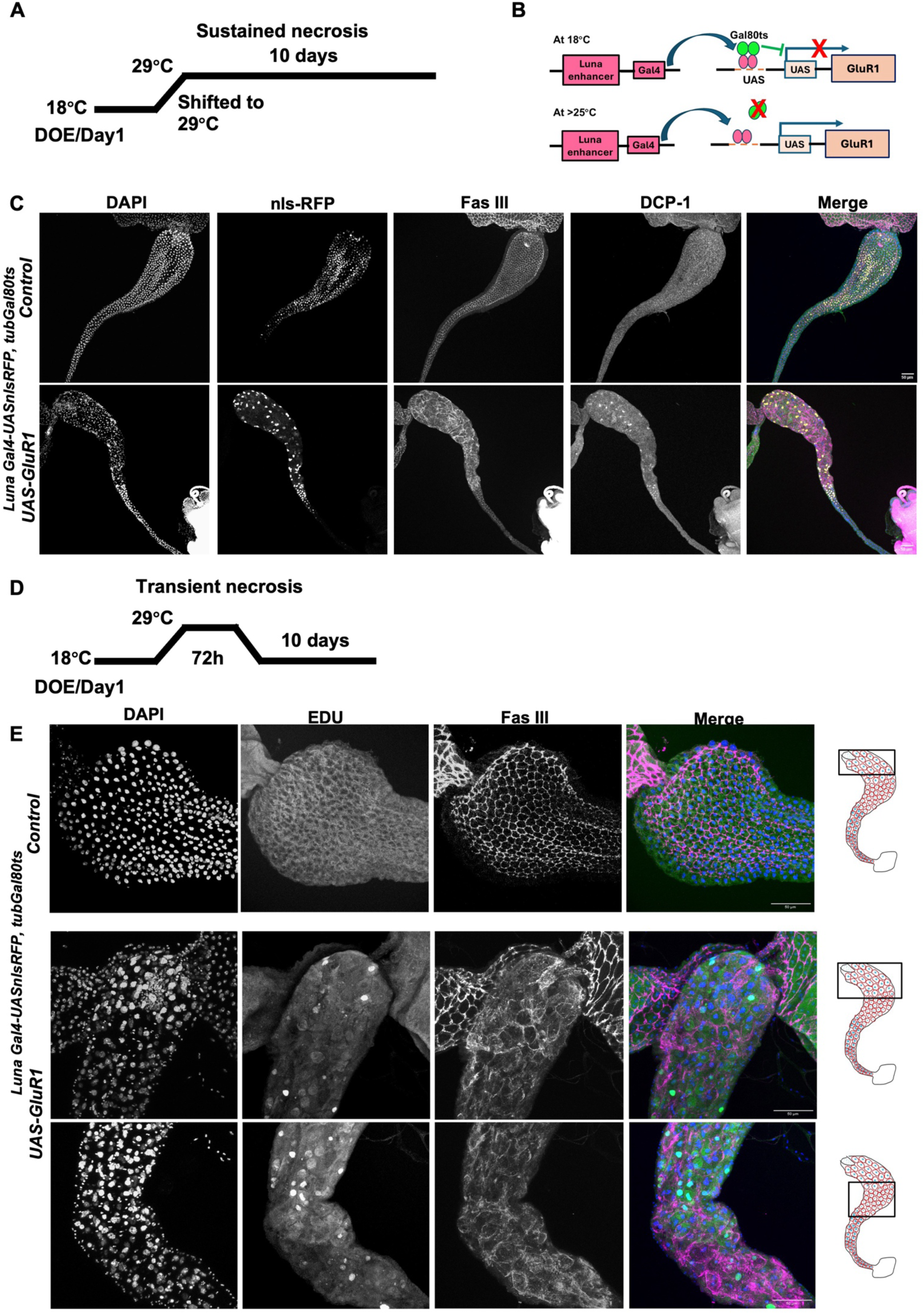
Transient necrotic damage extends the window of endocycling in the ED. (A,B) Schematic of GluR1 induction for sustained necrosis. (C) Confocal images of the entire ED in controls (top) or after 10 days of GluR1 expression at 29°C (bottom) stained with FasIII, DCP1 and DAPI. Control tissues express Luna-Gal4, UAS-RFP at 29°C without UAS-GluR1. (D) Schematic for transient GluR1 induction. Flies housed at 18°C suppress Gal4 activity via Gal80^ts^. On the day of eclosion, flies were shifted to 29°C for 72hours to induce GluR1 expression. Flies were returned to 18°C for 10 days to allow recovery. Controls experience the same temperature shifts but lack the GluR1 transgene. (E) Representative confocal images of the anterior ED region in controls (top) or the indicated regions of the ED transiently expressing GluR1 for 72h followed by 10 days of recovery in the presence of EdU. Note that the same ED is shown for the anterior and more posterior region of the GluR1-damaged sample, with some intentional overlap as indicated by the schematic at right.

We next shifted *Luna-Gal4/UAS-ReaperGal80^ts^* animals from 18 °C to 29 °C on the day of eclosion (DOE) for various timepoints to induce transient damage and allow for subsequent recovery at 18°C. After 32 hours of GluR1 induction we observed minimal organ damage, but after 72 hours of GluR1 expression, we observed reproducible cell loss in the absence of significant Dcp1 positivity (Fig. 5 D). We next examined ED organs after 10 days of recovery and observed significantly enlarged cells and nuclei (Fig. 5 E). To test whether necrosis induced cell loss engaged the endocycle for compensatory cellular hypertrophy, we induced tissue damage with GluR1 expression for 72h followed by immediate feeding with EdU during a 10d recovery period at 18°C. We observed robust EdU labeling in cells throughout the ED secretory epithelium, suggesting the longer damage paradigm with GluR1 engaged CCH in the anterior and posterior regions of the ED (Fig. 5E). The robust EdU labeling after 72h of damage also indicates necrosis induced cell loss extends the window of endocycling in the secretory epithelium, even beyond that of apoptosis induced cell loss.

Compensatory cellular hypertrophy can be triggered by a local alteration of tissue tension which accelerates the endocycle (Tamori and Deng, 2013). Here we show that the adult ED responds to a threshold level of cell loss to prolong the window of cell cycle plasticity in the adult to enable CCH to improve regenerative capacity. Future work will examine the signaling that determines the window of cell cycle plasticity in the adult ED and how cell loss and damage can extend this window.

## Materials and methods

### Drosophila husbandry

All the flies for constitutive Reaper expression for experiments in figure 1 and supplementary Figure 1 were housed at room temperature (23°C) on a Bloomington Cornmeal food recipe without crowding (<50 animals per vial). For the damage and recovery paradigm with Gal80^ts^, flies were housed at 18°C in the same media and shifted to 29°C for 32hrs for the activation of UAS-Reaper and shifted back to 18°C for the recovery. All the male flies used in all the experiments are virgins except for those used in fertility assays. We used the fly lines Luna-Gal4 (BDSC#63746), UAS-Reaper (BDSC#5824), Luna-Gal4 recombined with UAS-*nlsRFP* (BDSC#8546), UAS-Destabilized GFP (provided by D. McKay Lab, UNC Chapel Hill), GTRACE (BDSC#28280) and *Tubulin*-Gal80^ts^ (BDSC#7017), UAS-Hid (BDSC#65408), UAS CycE,UAS Cdk2 (Buttitta and Edgar, 2007), UAS-Dacapo-GFP (BDSC#83337), UAS-Stabilized Geminin (BDSC#59069), UAS-GluR1 (Klemm et al., 2021).

### ED dissection, fixation, and Immunostaining

*Drosophila* male EDs were dissected in 1X Phosphate buffered Saline (PBS), the dissected EDs were immediately transferred to an Eppendorf containing 1ml of 4% Paraformaldehyde (PFA) in 1X PBS to fix the tissues for 30 minutes under continuous rocking. Fixed EDs were rinsed twice using 0.1% Triton-X +1× PBS for 10 minutes each wash. Fixed EDs were permeabilized with 1% Triton-X+1X PBS for 30 minutes on a rocker. Permeabilized tissues were blocked with PAT (1X PBS, 0.1% Triton X-100, 1% BSA) for 10 min. Tissues were resuspended in primary antibody freshly diluted in PAT at an appropriate concentration (below) and incubated overnight at 4°C on a rocker. Tissues were then washed these tissues with 1XPBS 0.1% Triton X-100, 3 times 10 min. prior to incubation with secondary antibody in PBS-T (1× PBS + 0.1% bovine serum albumin + 0.3% Triton-X (PBT)-X, + 2% Normal Goat Serum (NGS)) for 4h at room temperature. Secondary antibodies were diluted to 1:2000. EDs were washed thrice with 1X PBS 0.1% Triton-X after the secondary antibody incubation for 10 minutes each wash. For actin staining we added 2 drops of Actin Red 555 in 1ml of PBS and incubated the EDs for 30 mins and washed twice with 10 mins each wash to remove unbound stain and incubated the EDs in DAPI 1µg/ml for 15 minutes to stain the DNA. EDs were washed with 1× PBS + 0.1% Triton-X after the DAPI staining and mounted in vectashield (Vector Labs). Tissues were mounted by creating a spacer between the slides and coverslip with double-sided tape or one layer of clear nail polish. The list of antibodies used is in the following table with their dilutions.

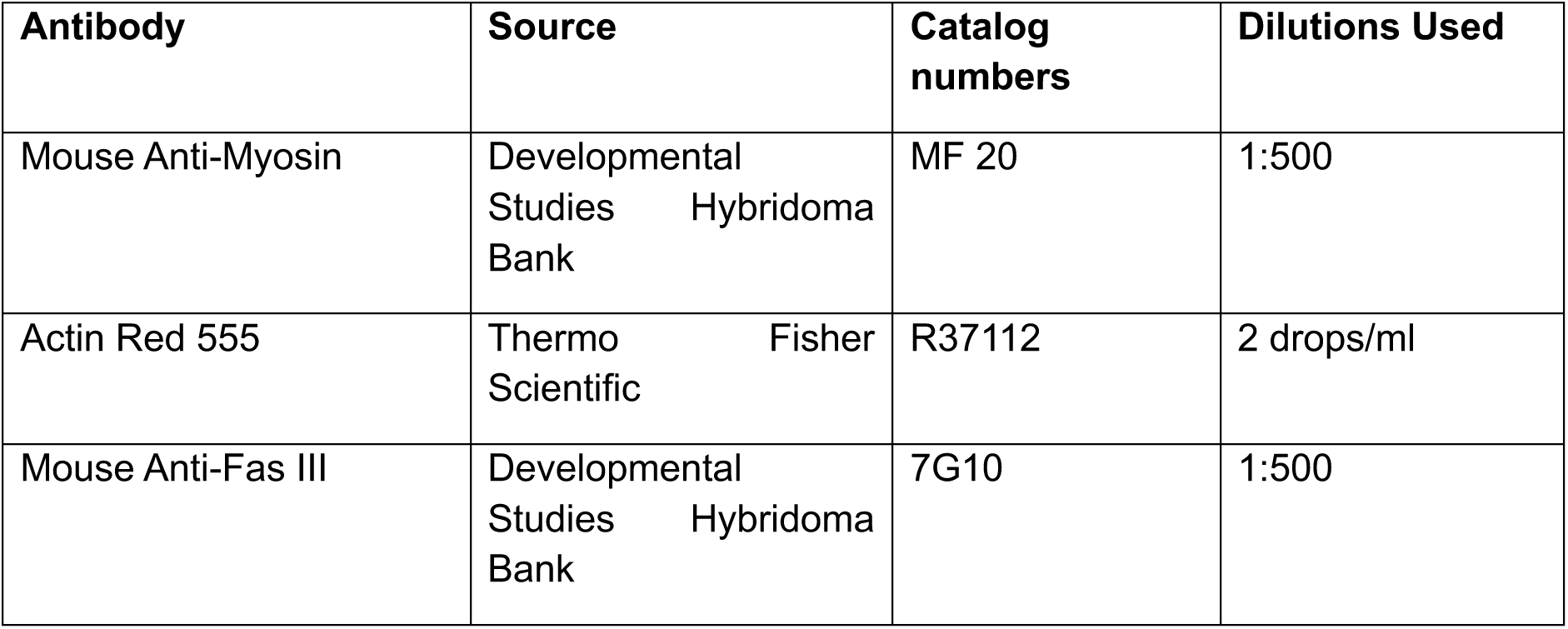
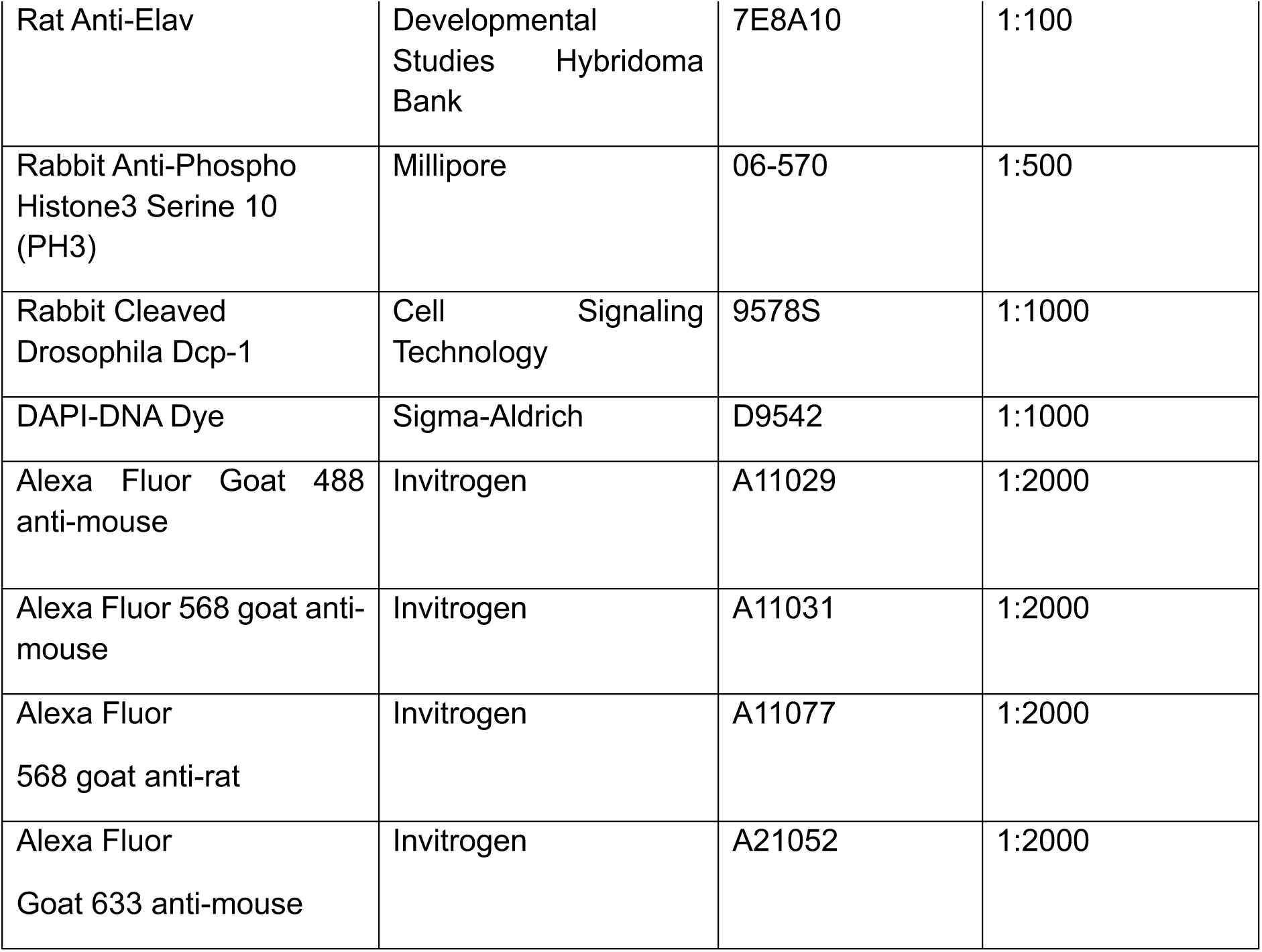

### 5-Ethynyl-2-deoxyuridine (EdU) feeding and labeling

To capture the endoreplication in ED secretory cells after inducing the damage, we fed the animals immediately after the 32hrs of 29°C damage (or 72h in the case of GluR1-induced damage) with an EdU solution made with 1mM EdU, 10% Sucrose, and blue food coloring (5µl/ml). We visually verified uptake of the EdU by the presence of food color in the abdomen. In cornmeal agar food vial, a Whatman paper was wetted with EdU solution (250µl in each vial). Flies were allowed to feed on this solution for 10 days during the recovery. The Whatman paper was replaced with a new one with a freshly made EdU solution every 2 days to prevent contamination due to the presence of sucrose. EDs were dissected at different time points during recovery when fed on EdU and proceeded with fixing, permeabilizing, and Click-IT reaction. We used Click-IT Plus EdU AlexaFluor-555 or 488 (Thermo Fisher C10638). We quantified the total number of EdU positive secretory cells by using the cell counter plugin in ImageJ Fiji.

### Confocal imaging

EDs for all the experiments were imaged using a Leica Stellaris or SP8 Confocal microscope with a 40X objective. Lower magnification 10X and 20X images were imaged using Leica SP5 Confocal microscope and Leica DMI6000B epifluorescence system.

### Area measurements

For the ED area measurements ED images were captured in one frame using a Leica SP5 Confocal microscope with 10X objective. The images were opened in Fiji Imagej software and obtained the maximum Z projection of the stacks of the image. Manually using the free hand selection tool, the area around the ED was drawn from the Anterior end of the ED where the AG, Seminal vesicles are connected to the posterior end of the ED where it connects to the EB. Area around the outlined ED was measured by clicking Analyze and then Measure. These area values were plotted using the Graph pad Prism. Incomplete or damaged ducts (due to dissection or mounting) were excluded from analysis.

For measuring the ED secretory cell sizes, in Fiji ImageJ we drew the area around the Fas III cell-cell junction staining of each cell using the free hand selection tool in the anterior region of ED and measured the area of each cell. For measuring the area of nuclei of secretory cells in the anterior region we took the maximum Z-projections of DAPI staining images and measured area for nuclei in the anterior region of ED.

### Ploidy/DAPI intensity measurements

All the ED DAPI images were obtained without saturating the DAPI using a Leica SP8 confocal microscope. Images were exported in Tiff format and opened in Fiji ImageJ. The maximum projection of the stacks of DAPI-stained images was obtained. We have observed that the basally located nuclei in the external muscle layer of the ED are diploid by flow cytometry. We used these diploid muscle nuclei of ED to normalize the measurements and to calculate the haploid nuclei intensity. We measured the background intensity by drawing the region of interest (ROI) in the background region of the images. After the background subtraction ROI was manually drawn around the DAPI-stained nuclei of Diploid and polyploid secretory cells. Measured the Raw integrated Intensity of both diploid and secretory cells polyploid nuclei. We measured around 30 secretory cell polyploid nuclei and 10 diploid nuclei from muscle for each ED and We calculated the corrected fluorescence intensity from each nucleus raw integrated intensity, by subtracting the average background intensity. The intensity of the haploid nuclei DNA was calculated by taking the average of the diploid nuclei intensity and dividing it by half, we used this as a reference of haploid nuclei with DNA content of 1C. Based on these haploid nuclei we estimated the DNA content of polyploid secretory cell nuclei. For Figures 4F we directly scored the DNA content without binning. For all other DNA content/ploidy measurements we plotted the after the following binning-2N (1.9-2.9), 4N (3.0-6.9), 8N (7-12.9), 16N (13.0-24.9), 32N (>24.9).

### Fertility Assay

Virgin males were used to set up the fertility assay in all the mentioned figures. For the fertility assay showed in figure 1 we used 2-3 day old virgin males expressing reaper and control were crossed to Canton S females. We used a ratio of 1:8 (Male: Female) to set up the fertility assay in each vial and allowed them to mate for 3 days in 25°C incubator. The total number of the pupae and the offspring eclosed were counted and plotted as the result of fertility. For each genotype we performed the fertility assay with 15 animals in each trial and repeated it three trials. For the fertility assay with Gal80^ts^ Reaper presented in Figure 4 we performed the fertility assay at two different temperatures. (i)For the post damage fertility test after the animals of both control and Gal80^ts^ Reaper underwent 32hrs 29°C damage and they were mated with virgin *Canton S* females with a ratio of 1:8 (Male:Female) in each vial in 29°C incubator to keep the damage on and allowed to mate for six days where we flipped them to fresh vial every two days. The total number of pupae were counted and plotted. (ii) For Day 10 post recovery after damage we performed the fertility assay after the 10 days of the recovery. The males were crossed to virgin *Canton S* females with a ratio of 1:8 (Male:Female) in each vial and allowed to mate in 18°C incubator for 6 days with every two days flipping and finally we counted the total number of pupae.

### Flow cytometry

Nuclei from the ED were isolated post dissection in S2 media (Thermofisher #21720024) with 10%Fetal Bovine Serum (Cytiva # SH30396.02HI) and 1% Pen-strep antibiotic solution. We used 25-30 intact EDs for the nuclear extraction per sample. We briefly spun the tissues to settle, S2 media was removed, then 1ml of Lysis Buffer (10mM Tris HCl-pH 7.4, 10mM NaCl, 3mM MgCl_2_, 0.1% NP40) was added and the sample was transferred to a Dounce homogenizer placed on ice. To release the nuclei we homogenized the tissues with 20 strokes of loose Dounce pestle and allowed the tissue to rest on ice for 5 minutes. Following this we homogenized with tight Dounce pestle for 40 strokes. We triturated the tissue 15 times with a silanized fire-polished glass Pasteur pipette (BrainBits). This homogenate was filtered through 100 microns filter mesh, followed by 40 micron filter mesh to remove debris and unlysed tissue. To this homogenate 100µl of S2 media was added to stop the lysis reaction. This filtrate was centrifuged at 800 rcf for eight minutes at 4°C. Removed the supernatant and added 200µl of S2 media to the nuclei pellet with and incubated on ice for 5 minutes. To this additional 800µl of S2 media was added and centrifuged for eight minutes at 800 rcf, 4°C, supernatant was removed and the pellet was resuspended in 200µl S2 media with Vybrant DyeCycle Violet (Invitrogen #V35003) at a ratio of 1:100 and allowed to incubate on ice for 5 minutes, after the incubation to this 800µl of S2 media was added and centrifuged at 800rcf for eight minutes at 4°C. The supernatant was removed and resuspended the pellet in 1ml of S2 media, gently vortexed, and run on an Attune NxT Flow Cytometer.

### Software and data analysis

We used Fiji ImageJ to analyze all the data. We used GraphPad Prism software to perform statistical significance in each graph plotted in the figures. The tests used to run the significance are inidcated in the figure legends.

## Acknowledgements

We thank Dr. Allison Box and members of the Buttitta Lab for helpful discussions regarding this project. We also thank Dr. Jill Haenfler and the lab of Dr. Josie Clowney for assistance with male courtship assays. N.R. was supported in part by an U. Michigan MCDB Edwards fellowship. Research reported in this publication was supported by the National Institute of General Medical Sciences of the National Institutes of Health under Award Number R35GM149273. The content is solely the responsibility of the authors and does not necessarily represent the official views of the National Institutes of Health. Stocks obtained from the Bloomington Drosophila Stock Center (NIH P40OD018537) were used in this study. Several antibodies used in this study were obtained from the Developmental Studies Hybridoma Bank (DSHB), created by the NICHD of the NIH and maintained at The University of Iowa, Department of Biology, Iowa City, IA 52242. We used FlyBase (most recently release FB2025_02, released April 17, 2025) to find information on Drosophila gene sequences, phenotypes, and gene functions (Ozturk-Colak et al., 2024).

## Notes

### Competing Interest Statement

The authors have declared no competing interest.

### Summary of Updates

Additional supporting data provided, including necrosis induced damage.

