## Supplemental Figures 1-5 for "A window of cell cycle plasticity enables imperfect regeneration of an adult postmitotic organ in *Drosophila*"

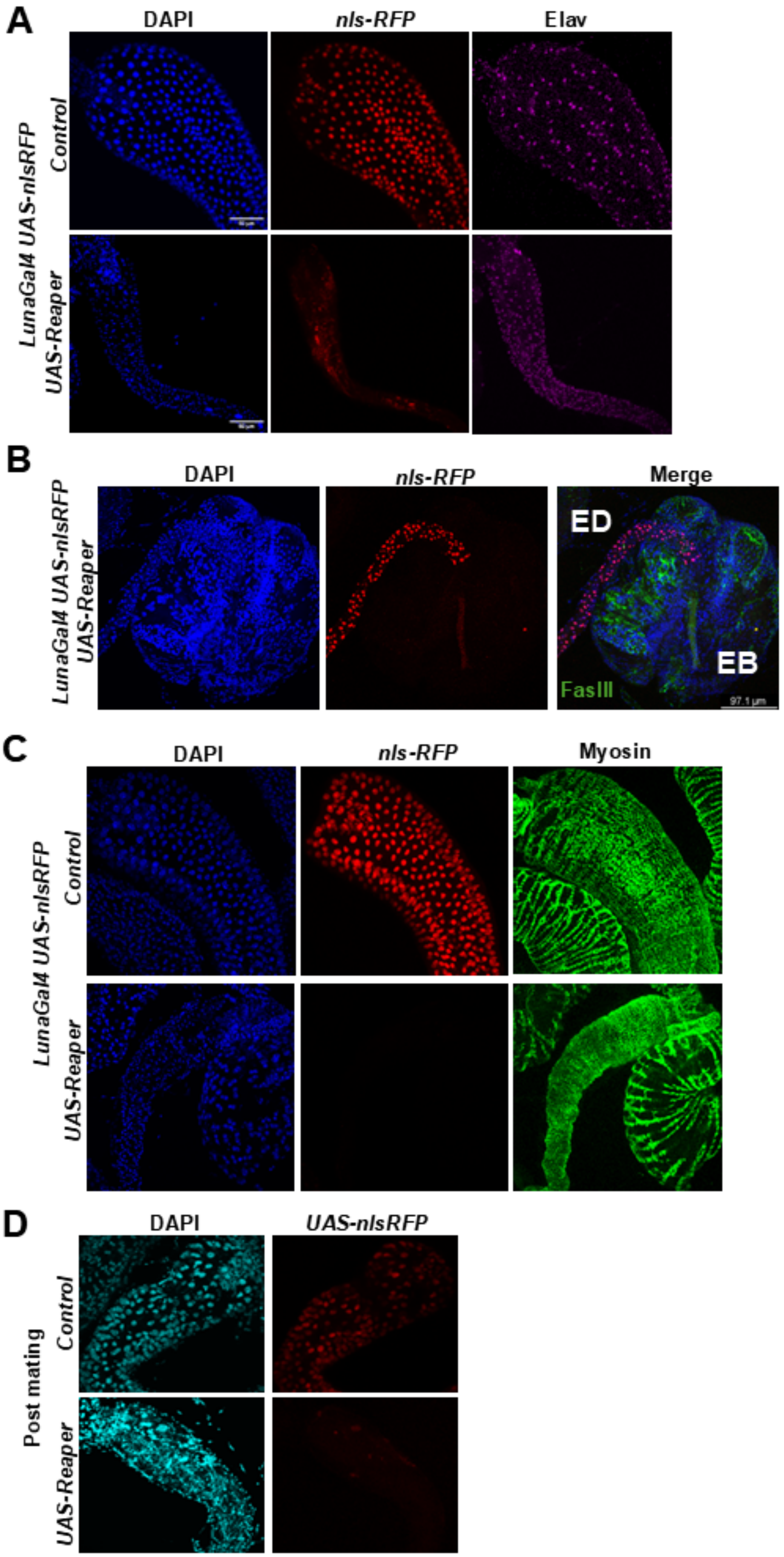
A
DAPI
nls-RFP
Elav
LunaGal4 UAS-nlsRFP
Control
UAS-Reaper
B
DAPI
nls-RFP
Merge
LunaGal4 UAS-nlsRFP
UAS-Reaper
ED
EB
FasIII
97.1 µm
C
DAPI
nls-RFP
Myosin
LunaGal4 UAS-nlsRFP
Control
UAS-Reaper
D
DAPI
UAS-nlsRFP
Post mating
Control
UAS-Reaper

**Supplementary Figure 1:**

(A) Confocal images of EDs constitutively expressing Reaper and UAS-nls-RFP in secretory cells. To validate Gal4 activity, nls-RFP (Red) was expressed under UAS control. Tissues were immunostained with Elav (Magenta), a neuronal/muscle nuclear marker. In Reaper-expressing EDs, complete ablation of secretory cells is evident, with only Elav-positive nuclei (neuronal/muscle) remaining, confirming selective loss of secretory cells.

(B) Confocal images of the posterior ED and ejaculatory bulb (EB) in control and Reaper-expressing flies. In Reaper EDs, nls-RFP–labeled secretory cells are observed being extruded apically through the EB.

(C) Confocal images of control and Reaper-expressing EDs stained with Myosin (Green) to label muscle fibers. Despite ablation of secretory cells, the muscle layer remains intact, indicating the specificity of Reaper-induced apoptosis to the epithelial cell population.

(D) Confocal images of the ED post-fertility assay in control and Reaper-expressing flies. In the Reaper EDs (lower panel), sperm accumulation is observed within the duct, suggesting impaired ejaculatory function due to secretory cell loss and tissue dysfunction.

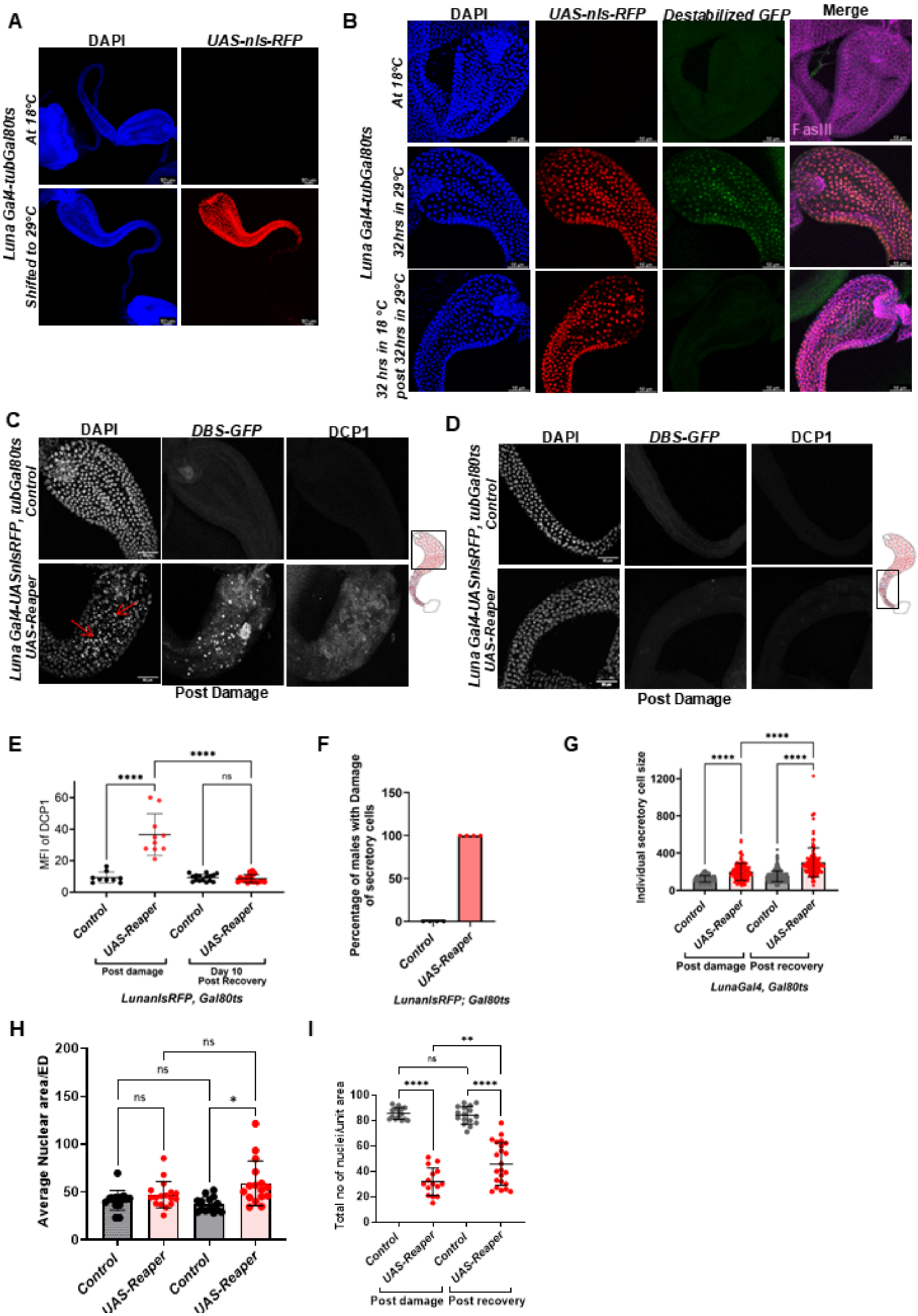
A
DAPI
UAS-nls-RFP
Luna Gal4-tubGal80ts
At 18°C
Shifted to 29°C
B
DAPI
UAS-nls-RFP
Destabilized GFP
Merge
At 18°C
32hrs in 29°C
32 hrs in 18 °C post 32hrs in 29°C
FasIII
C
DAPI
DBS-GFP
DCP1
Luna Gal4-UASnlsRFP, tubGal80ts
Control
UAS-Reaper
Post Damage
D
DAPI
DBS-GFP
DCP1
Luna Gal4-UASnlsRFP, tubGal80ts
Control
UAS-Reaper
Post Damage
E
MFI of DCP1
Control
UAS-Reaper
Control
UAS-Reaper
Post damage
Day 10 Post Recovery
LunanlsRFP, Gal80ts
F
Percentage of males with Damage of secretory cells
Control
UAS-Reaper
LunanlsRFP; Gal80ts
G
Individual secretory cell size
Control
UAS-Reaper
Control
UAS-Reaper
Post damage
Post recovery
LunaGal4, Gal80ts
H
Average Nuclear area/ED
Control
UAS-Reaper
Control
UAS-Reaper
I
Total no of nuclei/unit area
Control
UAS-Reaper
Control
UAS-Reaper
Post damage
Post recovery

**Supplementary Figure 2:**

(A) Confocal images of EDs showing temperature-dependent regulation of Luna-Gal4 by Gal80$^{ts}$. At 18°C (top panel), nls-RFP expression is suppressed, indicating Gal80$^{ts}$ effectively blocks Gal4 activity. Upon shifting to 29°C (bottom panel), nls-RFP is robustly expressed in the ED, confirming activation of Luna-Gal4 upon Gal80$^{ts}$ inactivation.

(B) Confocal images show *dsGFP* expression (reporting Luna-Gal4 activity) in EDs at different time points. At 18°C (top row), there is no *dsGFP* expression. After 32 hours at 29°C (middle row), *dsGFP* is detected in the anterior region of ED, confirming *Gal4* activation. Upon returning to 18°C, *dsGFP* signal disappears within 24 hours (bottom row), validating the kinetics of Gal80$^{ts}$-mediated repression.

(C) Confocal images of anterior EDs from control and Reaper-expressing flies after 32 hours at 29°C. In controls (top row), there is no signal from the *DBS-GFP* sensor or DCP-1 staining. In Reaper EDs (bottom row), strong *DBS-GFP* fluorescence and DCP-1 signal are detected, confirming caspase-mediated apoptosis in the anterior region of secretory cells.

(D) Confocal images of posterior region of ED from control and Reaper-expressing animals after 32 hours at 29°C. Both DBS-GFP and DCP-1 signals are absent in this region, demonstrating that Reaper-induced cell death is localized to the anterior ED.

(E) Quantification of mean fluorescence intensity of DCP-1 staining in EDs immediately after damage and after 10 days of recovery. Damaged EDs show significantly increased DCP-1 levels compared to controls, while no signal is detected after recovery.

(F) Quantification of the percentage of EDs that underwent secretory cell ablation following Reaper induction. Based on the presence of pyknotic nuclei, reduced ED size, and reduced secretory cell number per unit area, 100% of EDs exhibited damage.

(G) Quantification of individual secretory cell area from the anterior ED region in control, damaged (post 32-hour Reaper induction), and recovered (Day 10 post-damage) EDs. Each dot represents the area of a single secretory cell. Reaper-expressing EDs show a significant increase in cell size immediately after damage. Recovered EDs also display larger cell sizes compared to controls and damaged tissues, consistent with CCH.

(H) Average nuclear area per ED, calculated from the data in (i). Each dot represents the mean nuclear size per ED, supporting the observation of nuclear hypertrophy in damaged and recovered tissues.

(I) Quantification of the total number of nuclei per unit area in the anterior region of control and Reaper-expressing EDs, immediately after damage and post-Day 10 recovery. Reaper-expressing EDs show a reduction in nuclear density, consistent with cell loss and compensatory hypertrophy rather than proliferation during recovery.

Statistical analysis

(E) Descriptive percentage; 100% animals showed damage.

(E, G, H, I) One-way ANOVA with multiple comparisons between Day 0 and Day 10 for control vs. Reaper. P<0.0001 ****.

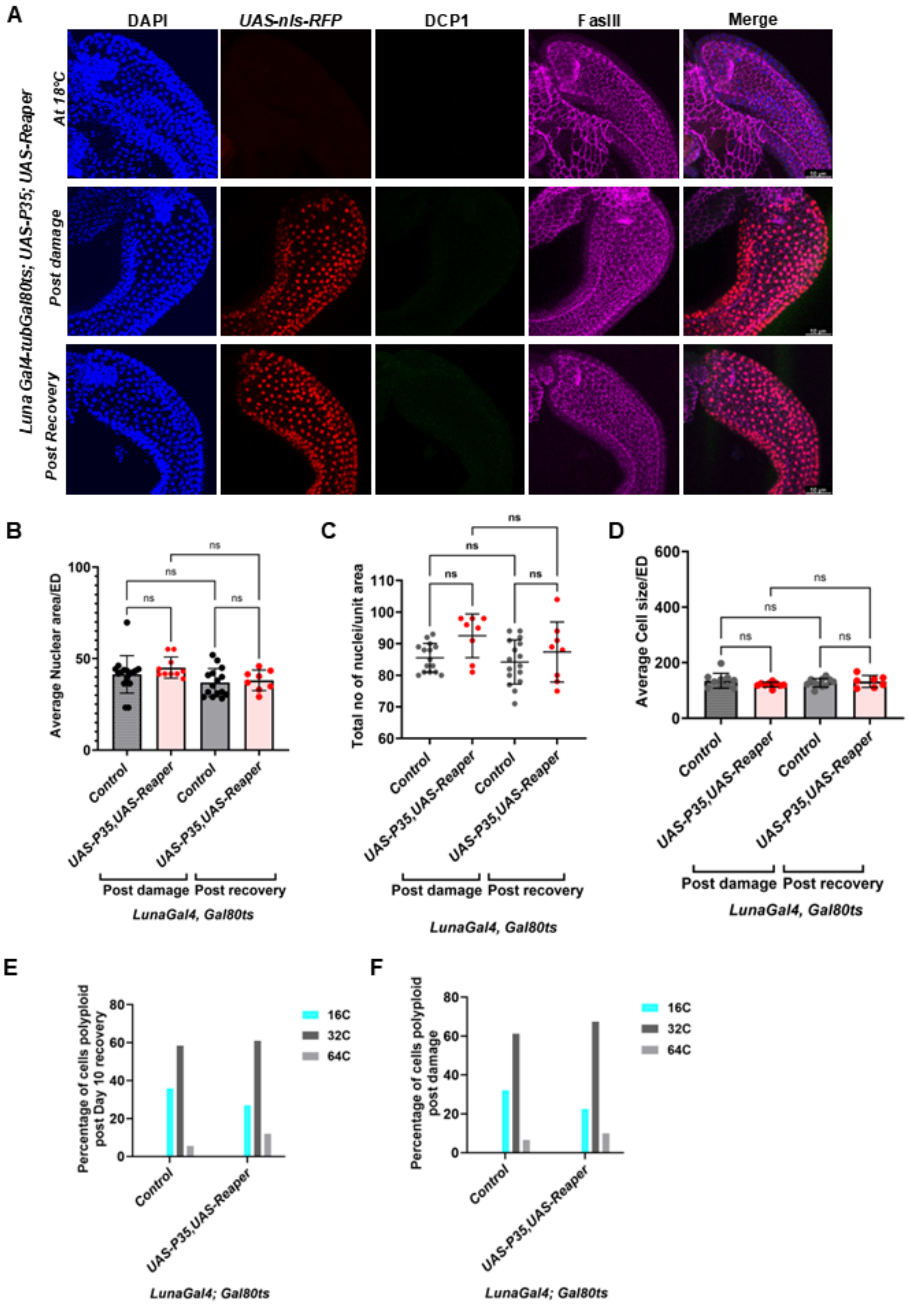
A
DAPI
UAS-nls-RFP
DCP1
FasIII
Merge
Luna Gal4-tubGal80ts; UAS-P35; UAS-Reaper
At 18°C
Post damage
Post Recovery
B
Average Nuclear area/ED
ns
Control
UAS-P35,UAS-Reaper
Post damage
Post recovery
LunaGal4, Gal80ts
C
Total no of nuclei/unit area
ns
Control
UAS-P35,UAS-Reaper
Post damage
Post recovery
LunaGal4, Gal80ts
D
Average Cell size/ED
ns
Control
UAS-P35,UAS-Reaper
Post damage
Post recovery
LunaGal4, Gal80ts
E
Percentage of cells polyploid post Day 10 recovery
16C
32C
64C
Control
UAS-P35,UAS-Reaper
LunaGal4; Gal80ts
F
Percentage of cells polyploid post damage
16C
32C
64C
Control
UAS-P35,UAS-Reaper
LunaGal4; Gal80ts

**Supplementary Figure 3:**
(A) Confocal images of ED with the transient expression of Reaper using $Gal80^{ts}$ showing the absence of damage to secretory cells with *UAS-P35* expression to block apoptosis. No damage to secretory cells in the ED is observed with Reaper + UAS-P35 induction for 32hrs at 29°C (Middle panel). After a Day 10 “post recovery” paradigm we did not see any differences in control vs. P35+Reaper expressing EDs. DCP1 staining was absent in the presence of UAS-P35 in ED.
(B) Quantification of nuclear area of secretory cells in control and UAS-P35 + Reaper-expressing EDs, immediately after 32hrs of damage paradigm and following Day 10 recovery paradigm. Each dot represents the average nuclear area per ED. We did not observe any significant increase in the nuclear area of controls without damage vs. UAS-P35 + Reaper expressing ED cells.
(C) Quantification of total nuclei per unit area in the anterior ED of control and UAS-P35+Reaper-expressing EDs both with the damage and recovery paradigm.
(D) Quantification of average cell size of secretory cells in control, UAS-P35 + Reaper-expressing EDs. There is no significant difference in the cell size of ED in both control and UAS-P35 + Reaper with both damage and recovery paradigm.
(E) Quantification of the ploidy of ED secretory cells in the anterior region of control EDs and UAS-P35 + Reaper immediately after the 32hrs damage paradigm.
(F) Quantification of the ploidy of ED secretory cells in the anterior region of control EDs and UAS-P35 + Reaper after the recovery paradigm.
Statistical analysis
(B, C, D) One-way ANOVA with multiple comparisons between Day 0 and Day 10 for control vs. Reaper. $P<0.0001$ ****.

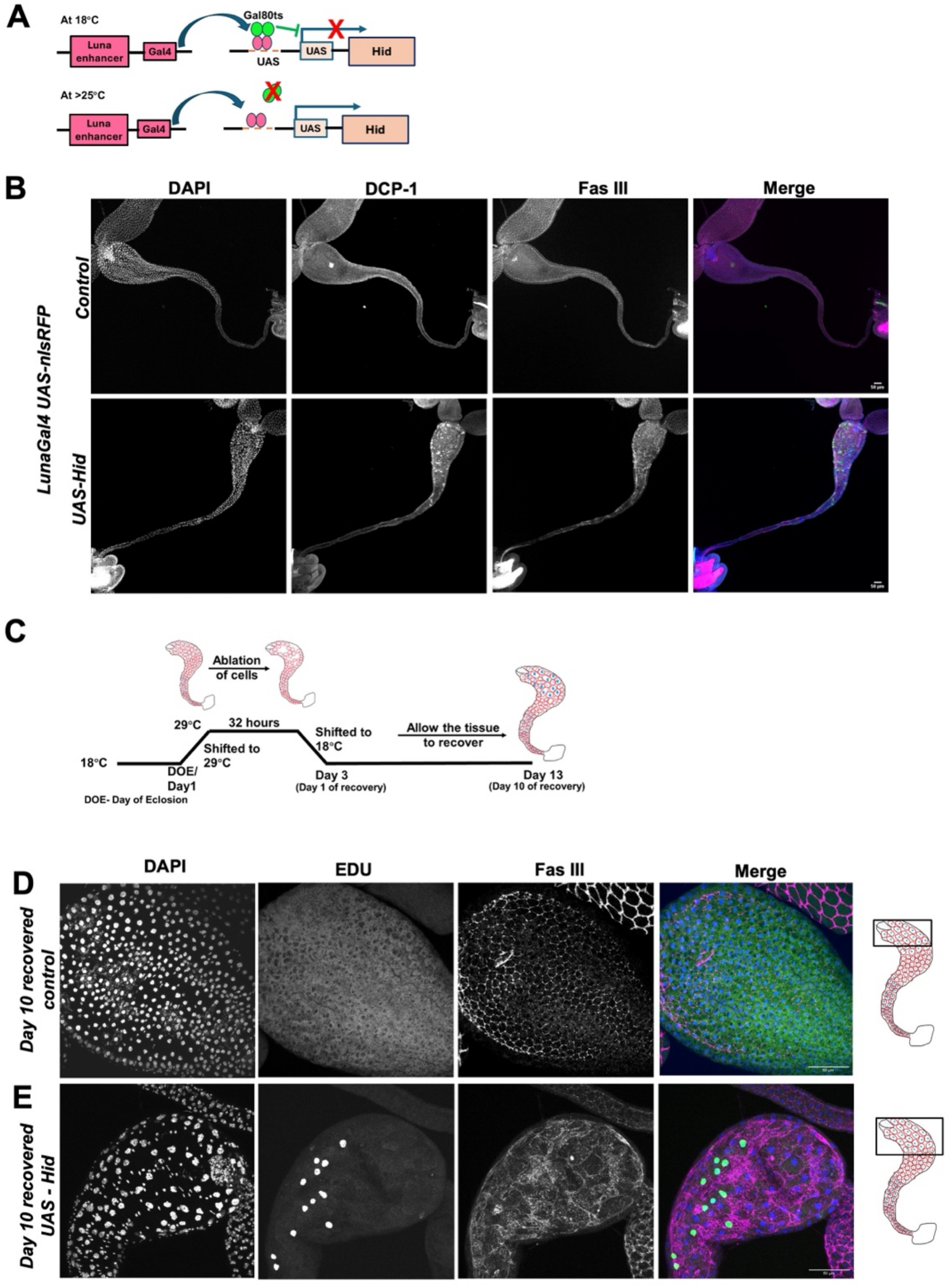


**Supplementary Figure 4:**

(A) Schematic showing targeted expression of Hid in the ED secretory epithelium.

(B) Confocal images of control vs Hid expression for 32h post-eclosion (PE), immunolabeled with FasIII, DCP-1 and DAPI.

(C) Schematic illustrating the use of the temperature-sensitive Gal80$^{ts}$ system to transiently induce Luna-Gal4-mediated expression of Hid in ED cells for 32h.

(D-E) Confocal images of the anterior region of ED immediately after 32 hours of Hid-induced damage induction followed by 10 days of recovery in the presence of EdU to label S-phases, FasIII to label cell-cell junctions and DAPI to label nuclei.

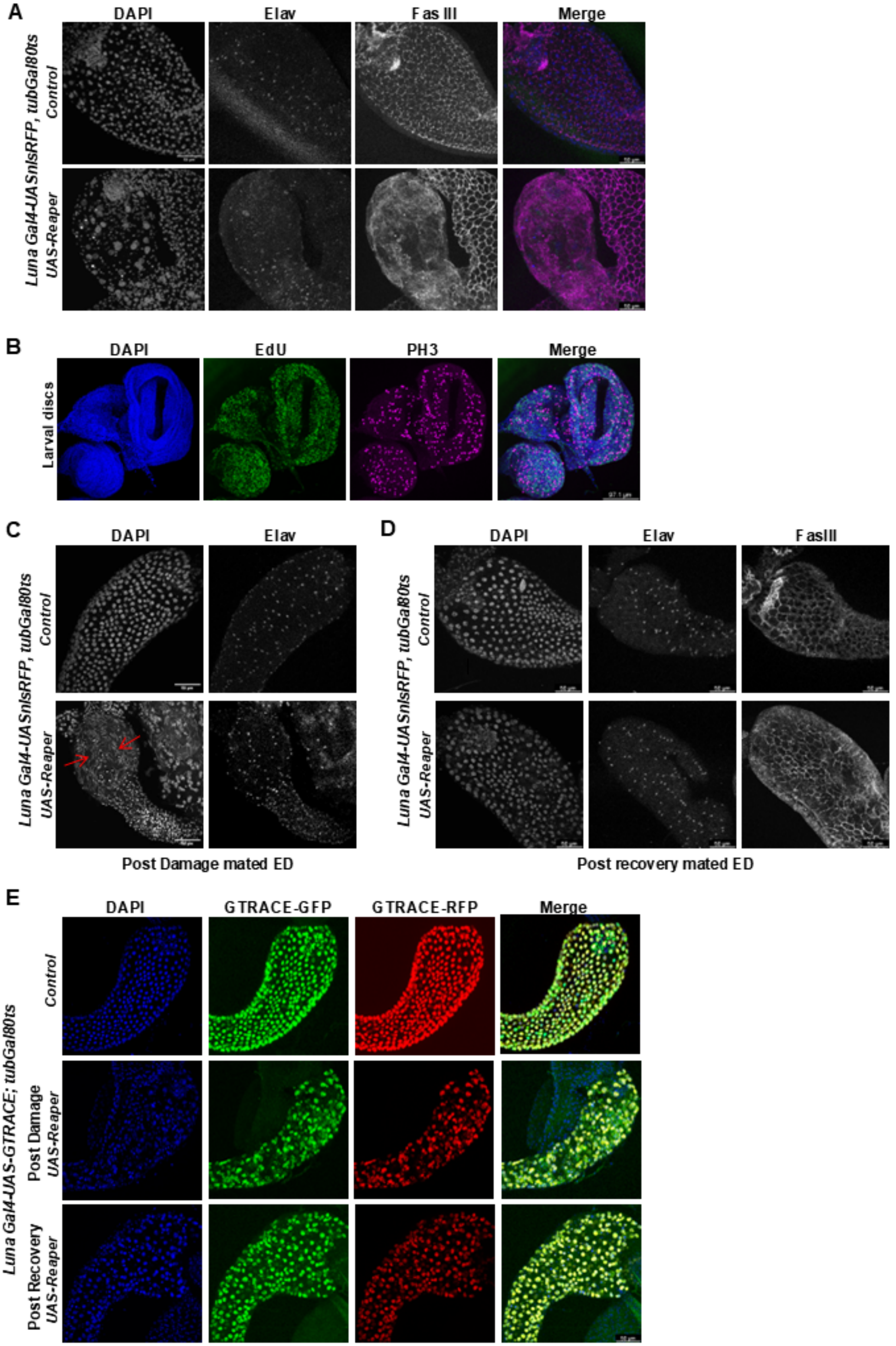
A
DAPI
Elav
Fas III
Merge
Luna Gal4-UASnlsRFP, tubGal80ts
Control
UAS-Reaper
B
DAPI
EdU
PH3
Merge
Larval discs
C
DAPI
Elav
Luna Gal4-UASnlsRFP, tubGal80ts
Control
UAS-Reaper
Post Damage mated ED
D
DAPI
Elav
FasIII
Luna Gal4-UASnlsRFP, tubGal80ts
Control
UAS-Reaper
Post recovery mated ED
E
DAPI
GTRACE-GFP
GTRACE-RFP
Merge
Luna Gal4-UAS-GTRACE; tubGal80ts
Control
Post Damage
UAS-Reaper
Post Recovery
UAS-Reaper

**Supplementary Figure 5:**

(A) Confocal images of anterior region ED stained with Elav, FasIII and DAPI of the control and the Reaper induced ED with the $Gal80^{ts}$ control system post recovery.

(B) Confocal images of larval imaginal discs stained with PH3 in red, EdU in green used as a positive control for the PH3 positive mitosis and nuclei stained by DAPI (blue).

(C) Confocal images of the ED post-fertility assay in control and Reaper-expressing flies immediately after 32hrs of damage. In the Reaper EDs (lower panel), sperm accumulation is observed within the duct, suggesting impaired ejaculatory function due to secretory cell loss and tissue dysfunction.

(D) Confocal images of the ED post-fertility assay in control and Reaper-expressing flies allowed to mate post day 10 recovery.

(E) Confocal images of ED expressing GTRACE in both control and Reaper under the control of $Gal80^{ts}$ immediately after damage and after 10 days of recovery. The *Luna-Gal4* driver used here is specific to the secretory cells of the ED in the male reproductive system and is active in the ED secretory cells from 1 day prior to eclosion throughout adulthood (Ramesh et al., 2025). We reasoned that if *Luna-Gal4* negative cells, possibly from the muscle layer or other nearby cell types, contribute to ED restoration through differentiation into the ED fate and subsequently turn on *Luna-Gal4* expression only after damage, their window of Gal4 expression prior to $Gal80^{TS}$ restoration would be very brief, resulting in temporally delayed and therefore limited expression of RFP (and possibly GFP) compared to resident ED cells that experience Gal4 activity immediately following the temperature-dependent inactivation of Gal80. This would be expected to result in much weaker expression of RFP, or possibly RFP “current” positivity without GFP “past” expression immediately after damage in non-resident derived cells. When we express G-TRACE under the control of *Luna-Gal4* with or without *UAS-Reaper*, we observe that all secretory cells in the ED co-express both GFP and RFP, immediately after damage as well as 10 days post-damage. These findings suggest that tissue mass recovery occurs from within the preexisting Luna-Gal4-expressing lineage of differentiated secretory cells, without the involvement of stem or progenitor cells.
